# Mind the Gap: An Embedding Guide to Safely Travel in Sequence Space

**DOI:** 10.1101/2025.06.19.660524

**Authors:** Adam Wu, Quentin Trolliet, Abhinav Rajendran, Jakub Lála, Stefano Angioletti-Uberti

**Author notes:** Electronic mail.

## Abstract

We present a hybrid approach combining a protein language model (pLM) with Monte Carlo (MC) sampling for generating enzyme mutants free of mutations deleterious for structural preservation. Given the amino acid sequence of the original enzyme and a set of residues for which the local environment should be conserved, i.e., the catalytic site, our approach generates mutants that differ vastly in the overall sequence while retaining the geometry of the conserved region, thereby representing promising candidates for further experimental screening. Unlike end-to-end deep learning approaches, whose results are harder to interpret and control, the use of a well-established, classic technique such as MC sampling allows us to easily interpret the generative process as the sampling of an energy landscape determined by the pLM. In turn, such an interpretation enables us to steer this generative process and control its outcome by making use of robust statistical mechanics concepts, e.g., temperature, thereby explicitly guaranteeing certain properties of the generated mutants. Given the increasing relevance of generative algorithms in the design and search for novel, optimised enzymes, we believe that our results constitute an important step for the future development of this class of techniques. To facilitate experimental verification, we finally provide hundreds of sequences for 13 different enzymes involved in catalytic processes ranging from carbon dioxide conversion to DNA replication.

## I. INTRODUCTION

Enzymes are protein catalysts that enable specific chemical reactions and are fundamental to numerous industrial and therapeutic applications, from biofuel production to pharmaceutical synthesis. Yet, the discovery and optimisation of enzymes with desired properties, such as enhanced activity, stability, or substrate specificity, remain a significant bottleneck in enzyme engineering.^1^ The current gold standard, directed evolution,^2,3^ is an iterative process that mimics natural selection by creating diverse libraries of enzyme variants through random mutagenesis or recombination, followed by high-throughput screening or selection for the desired trait. Although directed evolution has produced numerous successful enzymes, it is inherently limited by the vastness of sequence space.^1^ Experimentally probing this space is time-consuming and labour-intensive, and requires careful decisions about the initial sequence library. For example, one’s starting point might be enzymes from bacteria or plants that evolved in environments similar to those desired in the application. It is clear that in use cases with limited structural or mechanistic information, the challenge is even greater. One notable success story of directed evolution has been the discovery and optimisation of Taq-polymerase, a thermostable DNA polymerase that enabled the development of polymerase chain reaction (PCR) technology. Its discovery involved isolating the native enzyme from extremophilic bacteria that lived in the prohibitively hot springs of Yellowstone Park, where high temperatures caused the evolution of heat resistance necessary for PCR.^4^ Today, Taq polymerase is widely used for a wealth of biotechnology applications, from medical diagnostics to quality control in food production.^5^

When employing random mutagenesis strategies in directed evolution, a major limitation is often the high probability of introducing deleterious mutations: the intricate three-dimensional structure of an enzyme is crucial for its function and catalytic activity, and even a single amino acid substitution can disrupt this delicate architecture.^6^ Mutations that destabilise the structure of the active site are overwhelmingly likely to abolish or severely impair catalytic activity. Compounding this issue is the fact that functionally important residues are not solely confined to those close to the active site itself. Distant residues, both in terms of the primary sequence or of the three-dimensional structure, can still play a crucial role in determining the overall fold of the enzyme and thus maintaining the structural integrity of the active site. Therefore, a significant proportion of the mutants generated in directed evolution through random amino acid changes can lead to misfolding of the active site and loss of activity, representing wasted experimental effort and impeding the efficient exploration of functional enzyme space.

In recent years, the rapid advancement of artificial intelligence (AI) and the development of powerful protein Language Models (pLMs)^7–10^ have opened new avenues for protein engineering and enzyme design.^11–13^ In this regard, most AI-based generative approaches that appear in literature are one-shot generative approaches based on deep neural networks. While these models often differ in the architecture of the underlying neural network,^11–14^ the general idea underpinning the different models is similar: through training, these models learn the complex *joint* probability distribution of sequence and structure (or sequence-structure-function), and can use it to directly propose novel enzyme sequences with any desired characteristic. By learning these complex relationships, these models can, at least in theory, generate sequences that are more likely to fold correctly and possess the targeted activity without the need for extensive library construction and screening. Despite their promise, current end-to-end AI-based generative approaches often suffer from a critical limitation: the lack of interpretability. In other words, while these models can generate novel sequences exhibiting the desired properties, the inner workings of the generative mechanism and the underlying reasons for their success or failure, often remain opaque. This lack of interpretability hinders our ability to rationally refine the design process, troubleshoot unsuccessful designs, and extract generalisable principles for enzyme engineering. For example, two recently proposed algorithms for enzyme generation, ProGen^11^ and Raygun,^12^ lack the explicit possibility to preserve specific residues in the generative process, making it impossible to ensure the catalytic site remains present. In this case, multiple mutants must be generated and only those containing the catalytic site are chosen for downstream experimental testing. In general, understanding *why* a specific mutant is generated is crucial not only for building trust in these methods, but also for guiding future design efforts beyond a trial-and-error paradigm, and to integrate previous knowledge into the design campaign (for example, which residues are part of the catalytic site and must be preserved). These problems are further exacerbated if the goal is to build even more complex algorithms to drive multi-objective optimisation, where not only the catalytic activity but also other properties, such as thermal stability, should be enhanced.

To alleviate some of the aforementioned issues, we present a generative algorithm based on the relation between a specific residue and its embedding vector representation encoded by a pLM, describing its local context within the amino acid sequence. In practice, we use this relation to define an energy landscape where, by construction, the minimum energy corresponds to the target enzyme whose function we wish to mimic, and higher energies correspond to mutants with varying degrees of change in the environment around the residues that make up the catalytic site. By sampling this energy landscape via standard Monte Carlo (MC), we show how simply increasing the effective temperature allows us to reliably generate mutants that preferentially sample a larger neighbourhood around this minimum, while still remaining structurally close to the original enzyme and sample highly diverse sequences.

### 1. Using embedding similarity as a bias to generate mutants conserving the structure of specific regions

Decades of research on proteins and their evolution demonstrated that correlations between the chemical identities of amino acid residues can reveal how mutations propagate through a protein’s local environment.^7,15,16^ This correlation can be direct, e.g., when two residues are spatially close to each other, but also indirect, e.g., when they share a neighbour. As a result, when a single amino acid residue is altered, it can influence a neighbouring residue. This change then cascades, affecting other amino acids further along the chain, even if they weren’t directly adjacent to the initial modification. As such, rather than depending on their linear distance along the protein backbone, correlations are expected to decay with the distance between residues in the folded structure. This latter idea has been exploited to build protein folding algorithms, including the most recent and accurate ones, such as AlphaFold,^17^ RosettaFold^18^ or ESMFold.^7^ Among these algorithms, ESMFold derives these correlations between residues from a pLM called ESM-2,^7^ which was trained to solve a so-called masked-language problem. In this type of training, random residues are masked, and the algorithm is tasked with retrieving their identities given the surrounding sequence. Learning how to solve this problem is equivalent to learning correlations between residues: if the pLM can correctly identify that two residues in a sequence are strongly correlated, the presence of one of them can be used to increase the chance of correctly guessing the identity of the other, as opposed to a random guess. In technical, but quite illustrative jargon, these correlations are used to reduce the language model’s perplexity. Once trained, ESM-2 takes a sequence of amino acids as input and outputs contextual information in a multi-dimensional vector for each residue, its so-called embedding vector,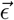. In a transformer architecture, embedding vectors are built as a combination of terms whose expansion coefficients are determined by the (self-) attention matrix 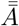. The entries of such matrix *A*_*ij*_ are interpreted as a measure of the strength of correlation between the (*i, j*) pair of residues, and, as a result, the embedding vector contains information regarding a residue within the context of the specific amino acid sequence provided as input. For this reason, these vectors can, and have been, successfully used as the starting input to a variety of algorithms for different downstream tasks. Crucially for our purposes, these tasks included structural ones, such as understanding the local fold to which a specific residue belongs to, or the prediction of residue-residue contacts. In fact, per-residue embedding vectors given by ESM-2 are used by highly accurate ESMFold to reconstruct a protein’s three-dimensional structure from its sequence alone.^7,19^

Following this interpretation, our main hypothesis is that if one changes the identity of one amino acid residue, the change in the embedding vector of the remaining ones is a measure of how much this mutation changed their local environment. Specifically, if a mutation in a residue *X* does *not* affect the local environment of a residue *Y*, then the embedding vector of residue *Y* should also be minimally affected, and vice versa. In this regard, we should also highlight that the magnitude of such a change should not only be affected by how strongly residues *X* and *Y* are correlated, but also by which amino acid *X* is substituted with. For example, substituting *X* with an amino acid that is biochemically similar might lead to a small change in *Y*, even when *X* and *Y* are strongly correlated. Importantly, pLMs encode knowledge of chemically conservative substitutions, allowing them to distinguish between structurally benign and disruptive mutations.

We formalise this previous hypothesis to build an energy function to quantify the effect of mutations. This energy function is then used within an algorithm that, starting from a reference protein, can generate mutants that retain the structure and local environment of any user-defined group of residues. Due to the structure-property relation, when this protein is more specifically an enzyme and the group of residues chosen is that of the putative catalytic site, we would expect such enzyme to also preserve its function. In other words, by preventing mutations that will make the enzyme unfold, our algorithm can generate a pool of candidates depleted in what would be otherwise deleterious mutations, thereby reducing the number of experimental screenings necessary in directed evolution approaches for enzyme optimisation.

A graphical representation of our algorithm is provided in Fig. 1. In simple words, our approach goes as follows: i) start with the original amino acid sequence of the enzyme to mimic; ii) propose a random mutation in one of the residues; iii) decide whether to accept or reject the proposed mutation based on how much the embedding vectors of the amino acids in the catalytic site have changed, using a Metropolis MC acceptance criterion; and iv) repeat points ii) and iii) to generate as many samples as requested. The general outline of this approach in itself is not novel. In the context of statistical sampling of a probability distribution, this algorithm has been used (e.g., for molecular simulations) at least since the 1950s,^20^ while also being recently employed in enzyme design.^21^ However, to the best of our knowledge, we are the first to propose the underlying acceptance criterion based on an energy function built on the connection between an amino acid sequence, its residues’ embedding vectors as provided by a pLM, and these residues’ local environment. This connection is crucial because it provides an easy route towards generating mutants that, while arbitrarily different from the original enzyme in any other aspect, can conserve any specific part of the enzyme and its structure. Unlike one-shot generative approaches based on deep-neural networks, where such conservation cannot be guaranteed, this capability allows us to generate mutants that preserve the key ingredients necessary to preserve catalytic activity. By avoiding deleterious mutations, our algorithm enriches the library in variants that are more likely to have higher fitness. Figure 2 illustrates this advantage and contrasts embedding-guided sampling with the purely sequence-based exploration typical of directed evolution. Additionally, the definition of a generalised energy to produce mutants via MC sampling allows us to provide a robust interpretation of the generative procedure in terms of statistical mechanics concepts.

**Figure 1.**
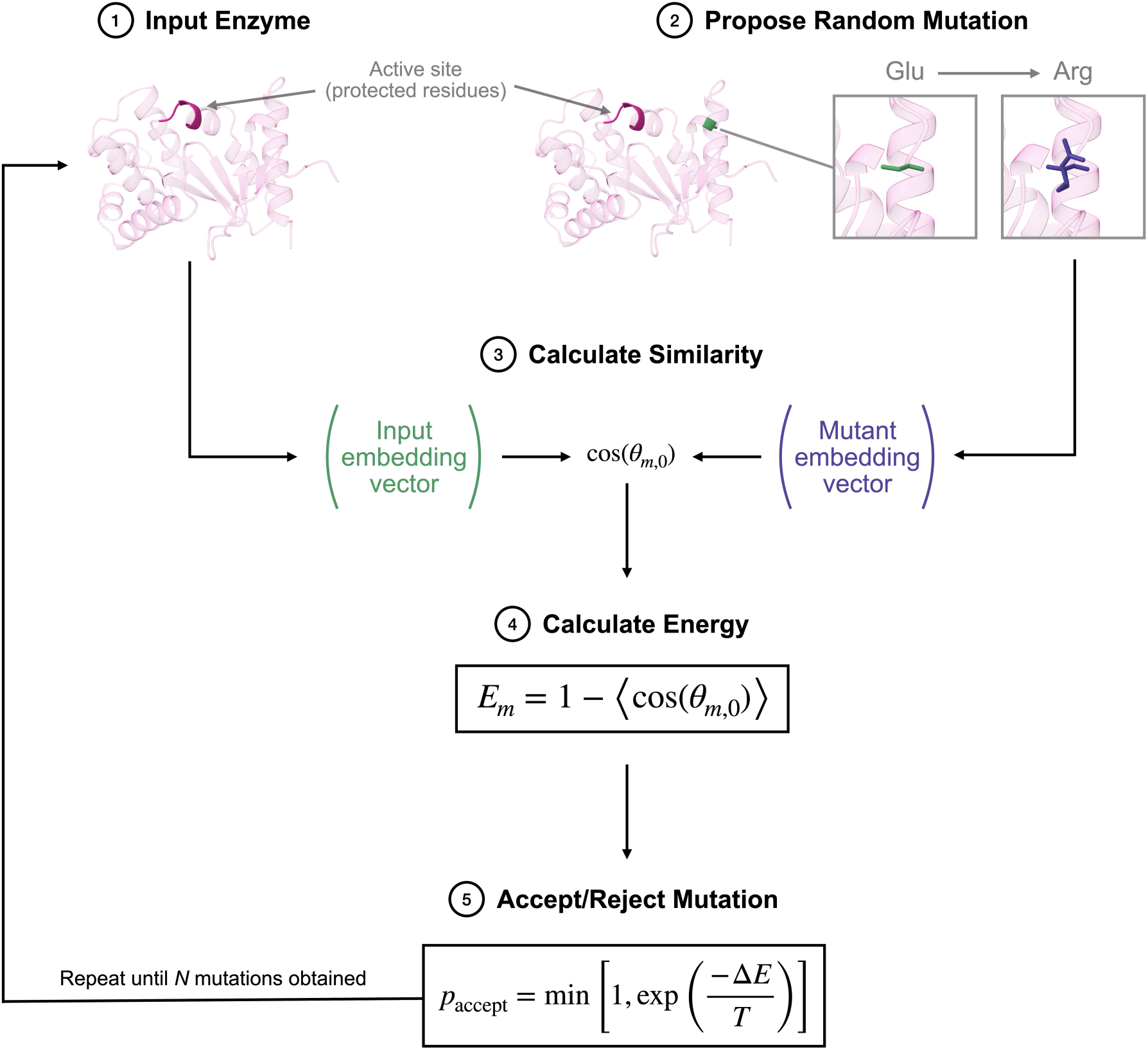
Flow chart summarising our generative algorithm. Starting from a reference enzyme as input, a random mutation is made. The impact of this mutation is quantified by computing the change in embedding vectors, derived from the ESM-2 pre-trained pLM, for a user-defined set of residues X whose local environment is to be preserved. This change, measured by cosine similarity, defines an effective energy used to guide a standard Metropolis MC sampling algorithm to accept or reject the mutation, following Eq. 1. The well-known mathematical properties of this type of algorithm guarantee that the equilibrium distribution of sequences generated spans all the possible mutants within a certain energy from the original one. Correspondingly, when the connection between a residue’s embedding vector and its environment holds, such a procedure generates mutant sequences that maintain the spatial configuration around *X*. If the choice of {*X*} carefully includes all residues in the catalytic site, this process effectively generates a library of enzyme variants with likely preserved catalytic activity.

**Figure 2.**
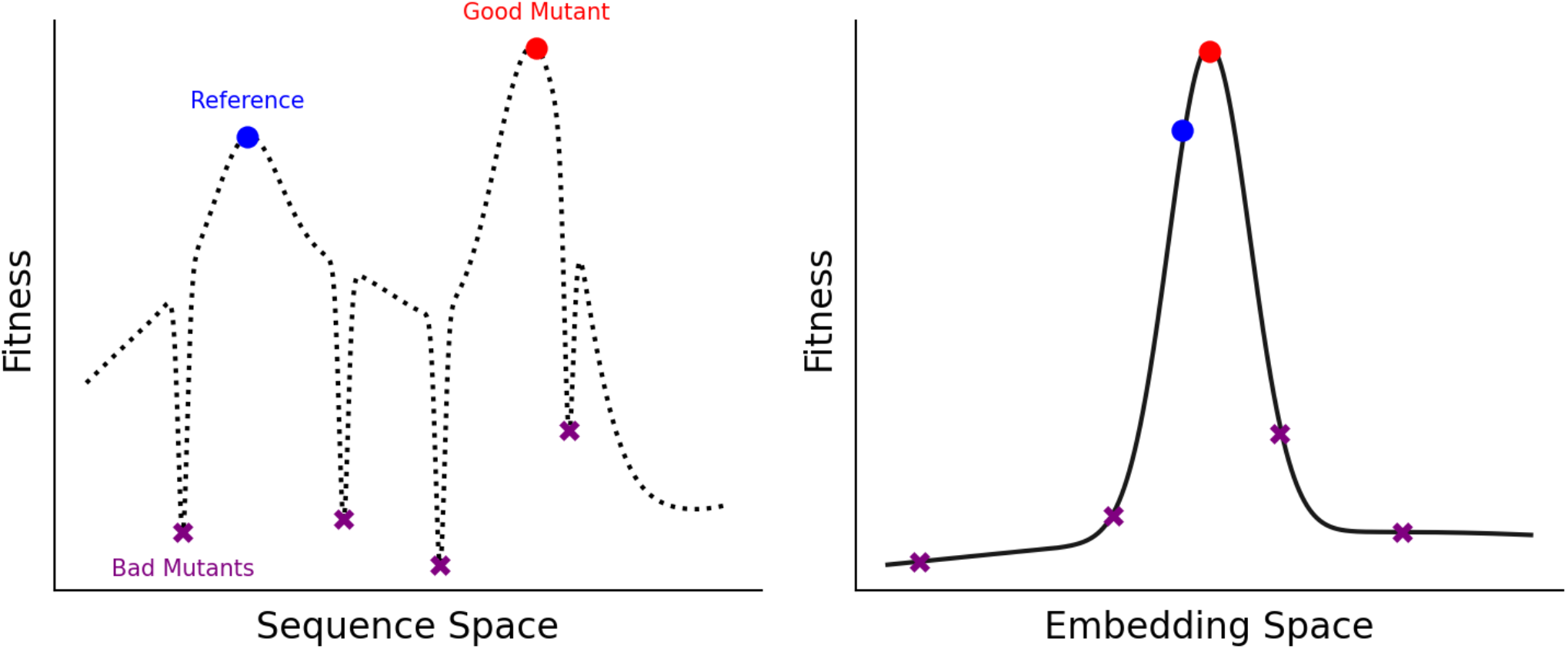
Fitness landscape schematic comparison between sequence and embedding spaces. In sequence space (left), structurally or functionally similar mutants, e.g., the reference enzyme and a viable mutant, are often separated by low-fitness intermediates, making naive exploration problematic and inefficient in directed evolution. In contrast, in embedding space (right), such mutants can cluster together, smoothing the landscape and removing fitness “gaps”. Sampling in the embedding space enables the generation of diverse, high-fitness candidates while avoiding deleterious variants, making it a more effective starting point for downstream experimental optimisation.

### 2. An energy function built on residues’ embedding similarity and its use for sampling

Before we analyse the results of our generative approach, we briefly outline the underlying mathematical framework and highlight key properties of the sampling algorithm. Central to our method is an energy function defined in the latent space of the ESM-2 pLM, which encodes contextual embeddings for each residue in a protein sequence. We define the energy of a mutant, *E*_*m*_, with respect to a reference sequence of amino acids, {*X*}, as:

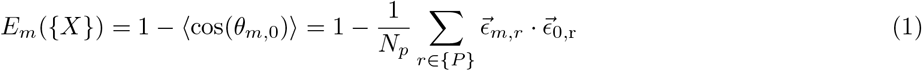

Here,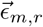is the embedding vector for residue *r* in mutant *m*, and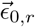 is the corresponding embedding in the reference sequence. The sum is taken over a designated subset of *N*_*p*_ residues that are part of the group {*P*} *ρ* {*X*}, which we refer to as the *protected* residues. Specifically, these are residues whose local environment will be preserved. In the case of an enzyme, the protected residues are typically those directly involved in the catalytic site, or those located within a certain distance from it in the folded structure. The energy function therefore quantifies the extent to which the embeddings, and by extension, the local environments, of the protected residues have changed in the mutant sequence. Importantly, *E*_*m*_ is not the inverse of a fitness function (as in Fig. 2) but is instead defined in an *effective* embedding-similarity space.

Using this definition of the energy, sequence generation proceeds via a standard Metropolis MC sampling procedure.^22^ At each step, a random mutation is proposed to any of the amino acids that are not part of the protected group, and is accepted with probability:

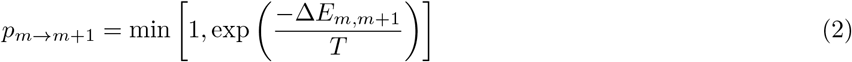

where t:i. *E*_*m,m*+1_ = *E*_*m*+1_ − *E*_*m*_ is the difference in energy between the proposed mutant at step *m* + 1 and the current state of the system *m*.

From a mathematical point of view, the chain of states (i.e., amino acid sequences) generated by using a Metropolis MC algorithm and the acceptance criteria defined by Eq. 2 is a discrete-time Markov Chain^23^ sampling a Boltzmann probability distribution characterised by the energy function *E*. Under very mild assumptions, statistical mechanics thus guarantees that our algorithm will sample a region minimising the free-energy *F* (*T*) = *E* − *TS*(*E*), where *S*(*E*) = *k*_*B*_ ln *W* (*E*) is the entropy of the latent space at a specific energy *E*, and *W* (*E*) is the number of states (i.e., amino acid sequences) with such energy.

This formulation presents us with an important property. The effective temperature *T* controls how far in energy mutants will sample from the energy minimum, which by construction, corresponds exactly to the reference sequence. In other words, starting from our reference sequence, and after an initial transient, the equilibrium distribution sampled by our generative algorithm will correspond to sequences within a certain distance in latent space from the reference sequence. If a small distance in energy or latent space corresponds to sequences whose folded structure preserves the local environment of the protected residues, our algorithm guarantees to generate similarly folded sequences, regardless of other differences. For example, while a low temperature will limit the distance sampled in latent space from the original enzyme, there is nothing limiting sequence diversity besides the explicit conservation of the protected residues imposed by the mutation proposal. Thus, the number of amino acids that are different in the reference compared to the mutant can be, at least in principle, arbitrarily large to the upper limit of *N* − *N*_*p*_, where *N* is the total sequence length. While correlations between sequence and structure will still implicitly limit how far in sequence the mutant can be from the reference enzyme, our algorithm is mathematically guaranteed to sample all sequences within a certain energy region without any bias, and thus generate the maximum diversity allowed within this space.

## II. RESULTS

In Fig. 3, we show twelve illustrative examples of mutants from four representative enzymes, generated using our approach, superimposed onto folded structures for the original enzyme. The folded structures from which the root mean square deviation (RMSD) was extracted have been calculated using ESMFold (see details in the Methods section). To reduce the possibility of having generated adversarial sequences that might mislead the protein folding algorithm, calculations using AlphaFold2 have also been performed. These results are provided in the Supplementary Information and fully confirm all trends observed using ESMFold.

**Figure 3.**
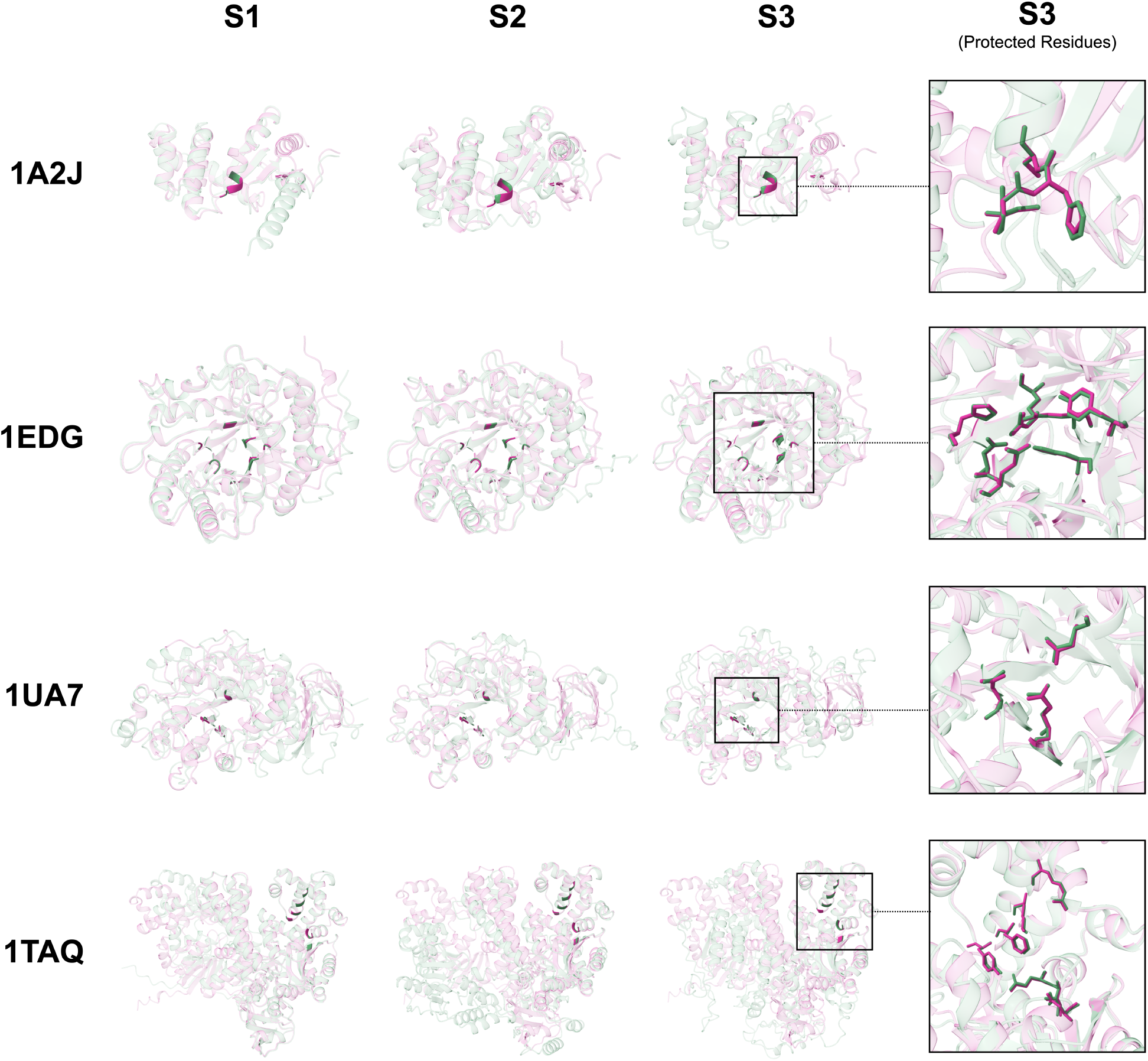
Examples of mutants for different representative enzymes generated using our methodology, highlighting its protected catalytic site. The pink-shaded cartoon is a ribbon representation of the structure of the original enzyme, and the green-shaded ones are the mutant (both predicted by ESMFold^7^), overlaid on top of each other. Dark regions correspond to the protected residues in the catalytic site. The panel on the right hand side magnifies the catalytic site of the enzyme. While the root mean square deviation (RMSD) between the catalytic site of the original enzyme and that of the mutant remains extremely small (less than 0.3Å), the difference in the rest of the structure can be almost arbitrarily large, demonstrating the ability of our algorithm to generate very diverse enzymes while preserving the geometry of specific, user-defined regions. Additional details for these mutants can be found in Table I. The PDB codes for the reference enzymes are: 1A2J = oxidoreductase, 1EDG = cellulase, 1UA7 = hydrolase, 1TAQ = Taq polymerase.

In Table I, we also report the RMSD of the protected residues between the original enzyme and the mutants. Across all reported structures, the RMSD remains below 0.3Å, indicating exceptional preservation of local geometry. This structural conservation is particularly remarkable given the broad range of sequence identities between mutants and the reference enzymes, which span from as low as 9% to as high as 95%. This proves that our algorithm is capable of generating mutants that, despite an almost identical environment for the protected residues, can differ largely in sequence.

**Table I.** Summary of the enzymes analysed in the main text along with their putative protected residues. For each enzyme, structural comparisons between three engineered mutants (S1, S2, S3) and the reference structure are quantified using the root-mean-square deviation (RMSD) of the protected residues. Sequence identity percentages for the full-length enzymes relative to the parent are also provided. Additional enzymes and alignment analyses are detailed in the Supplementary Information.

| PDB ID | Class | Protected Residues | Protected RMSD (Å) |  |  | Sequence Identity (%) |  |  |
| --- | --- | --- | --- | --- | --- | --- | --- | --- |
|  |  |  | S1 | S2 | S3 | S1 | S2 | S3 |
| 1A2J | Oxidoreductase | Cys-30, Cys-33 | 0.035 | 0.033 | 0.044 | 53 | 24 | 9 |
| 1EDG | Cellulase | Arg-79, His-122, Asn-169, Glu-170, His-254, Tyr-256, Glu-307 | 0.188 | 0.184 | 0.204 | 81 | 51 | 44 |
| 1UA7 | Hydrolase | Asp-173, Glu-205, Asp-266 | 0.243 | 0.070 | 0.229 | 39 | 29 | 14 |
| 1TAQ | Taq polymerase | Asp-610, Ile-614, Glu-615, Phe-667, Tyr-671, Lys-663, Arg-659 | 0.130 | 0.234 | 0.280 | 95 | 71 | 27 |

Although the protected residues exhibit minimal structural deviation, the remainder of the protein structure shows greater variability. As visualised in Fig. 3 and Supplementary Fig. S-3, RMSD values outside the protected site can vary substantially. In some cases, the ratio between the global RMSD and the RMSD of the protected residues (both calculated using the original enzyme as reference) approaches three orders of magnitude, ranging from approximately 1 to 400. This broad variation reflects a key design feature of our method: it does not impose constraints on structural regions beyond the specified protected residues, allowing for substantial backbone and side-chain flexibility elsewhere in the protein. We also notice that all predictions made by ESMFold have very high confidence. The average predicted Local Distance Difference Test (pLDDT),^24^ a standard metric of folding algorithm confidence in local structure predictions around a certain residue,^7,17^ is very high for protected residues, with a minimum of 0.87 for all reported structures in Fig. 3. In contrast, pLDDT scores for non-protected regions show greater variability, reflecting the trends observed in RMSD. Together, these results confirm that our embedding-guided algorithm reliably preserves the structural integrity of catalytically relevant regions, even when extensive sequence-level diversity is introduced.

### 1. Embedding similarity is indeed a good proxy for similarity in the local structure

While the results above qualitatively demonstrated that our generative approach can produce sequence-divergent mutants with preserved local structure at user-defined residues, we now quantitatively assess the relationship between our embedding-based energy function (Eq.1) and structural similarity, as measured by RMSD. This analysis is shown in Fig. 4, which reports the RMSD of protected residues compared to the corresponding energies for a large number of mutants derived from the representative enzymes shown in Fig. 3. A complementary analysis in Fig. 5 reports a similar graph, but with the average pLDDT instead of the RMSD of the protected residues (additional data is provided in the Supplementary Information).

**Figure 4.**
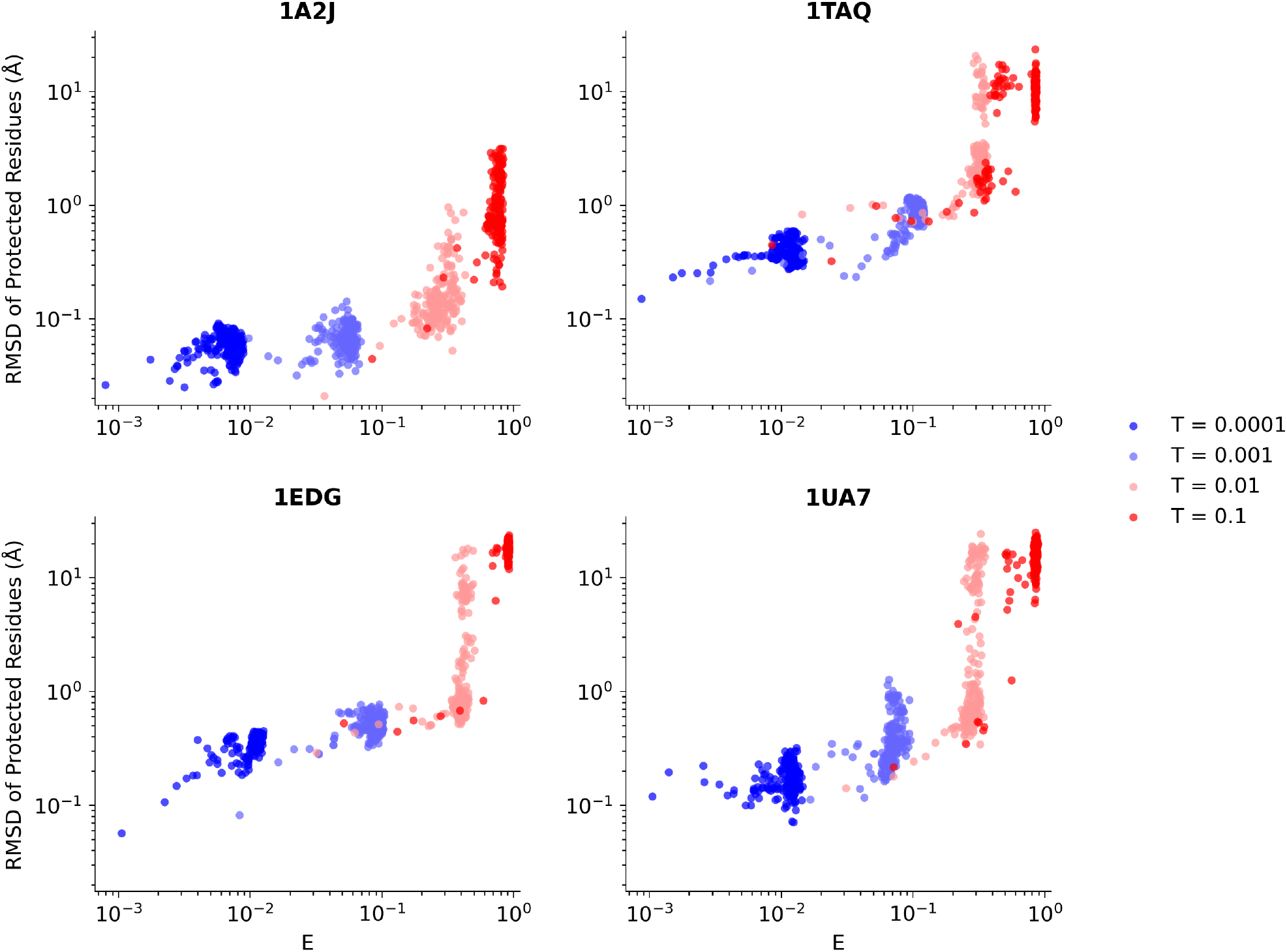
Scatter plots of the energy, *E*, vs. the RMSD of the protected residues, for the four representative enzymes from Fig. 3. We use the Spearman’s rank correlation coefficient, *ρ*, reported in Table II, to measure the monotonicity of the function describing the connection between two variables plotted here. There is a strong correlation between the quantities, with lower energy values corresponding to lower RMSD values for the protected residues, allowing one to use the former as a proxy for the latter during the sampling procedure.

**Figure 5.**
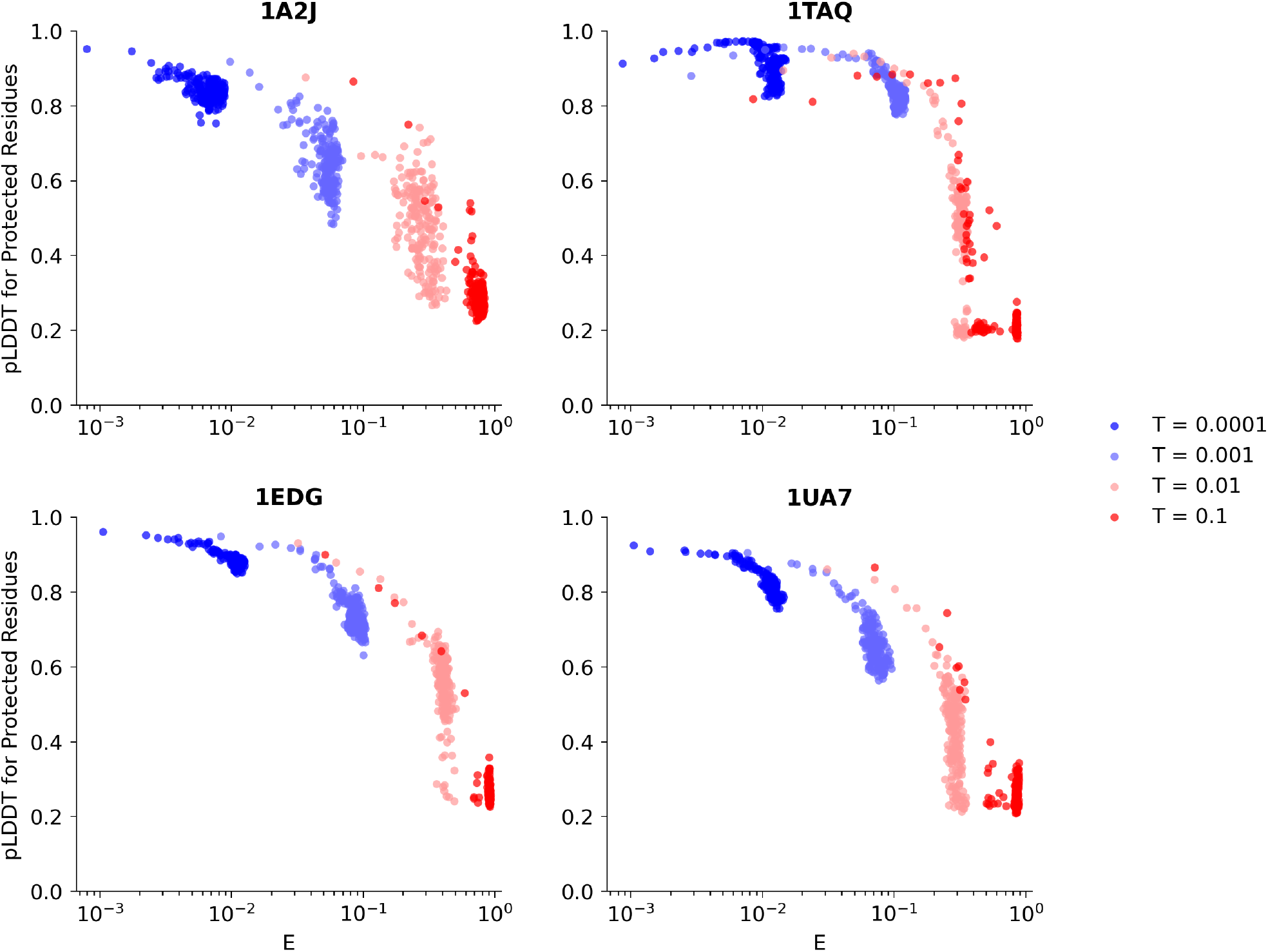
Relationship between pLDDT and mutation energy (*E*) for protected residues across four representative enzymes from Fig. 3. The scatter plots reveal a consistent negative correlation: residues with lower mutation energies tend to exhibit higher pLDDT scores in ESMFold predictions. This trend is most prominent at low sampling temperatures, particularly at *T* = 0.0001, where all pLDDT values exceed 0.75, suggesting satisfactory structural confidence. The strength of these correlations, quantified via the Spearman’s rank correlation coefficient, *ρ*, is detailed in Table II.

To quantify these trends, we calculate the Spearman’s rank correlation coefficient *ρ*,^25^ a statistical measure of a function’s monotonicity, whose modulus varies between 0 (no correlation) and 1 (perfectly monotonic function), with the sign indicating increasing (positive) or decreasing (negative) behaviour. As can be observed in Table II, there is a consistently strong positive correlation between embedding-based energy and RMSD (*ρ* ranging from 0.848 to 0.931), as well as a strong negative correlation with pLDDT (*ρ* ranging from –0.852 to –0.944). These results confirm that lower-energy mutants, i.e., those more similar to the reference in embedding space, also tend to be more structurally similar and confidently predicted.

**Table II.** Spearman’s rank correlation coefficient *ρ* between the mutant energy *E* (see Eq. 1) and the RMSD of protected residues in the mutants generated in this study. The strong correlation, signalling a monotonic relation between energy and RMSD, supports the interpretation of the embedding vector as a descriptor for the local residue environment. Notably, a similarly strong correlation, but of opposite sign, connects our energy to the pLDDT, with the reported values also signalling very high confidence in the predicted structures for low enough energies. All reported correlations have a p-value *<* 10^*-*4^.

| Enzyme (PDB) | $\rho(E, \text{RMSD})$ | $\rho(E, \text{pLDDT})$ |
| --- | --- | --- |
| 1A2J | 0.855 | -0.877 |
| 1EDG | 0.931 | -0.944 |
| 1UA7 | 0.848 | -0.852 |
| 1TAQ | 0.924 | -0.944 |

This is a critical finding, because while our MC procedure guarantees sampling over embedding-defined energy landscapes, this energy is only defined through the sequence embeddings of the protected residues, not structural deviations. Initially, we only hypothesised that these two quantities should have a strong correlation; here, we provide empirical evidence for this assumption (at least in a statistical sense). From a practical perspective, the observed correlation means that we do not need to fold structures at each step during sampling, which would increase computational costs by orders of magnitude. Instead we can simply take changes in the embeddings as a useful, computationally cheaper proxy to drive the generative procedure. In other words, to generate mutants with a conserved environment for the protected residues, we only need to to sample low-energy regions in the embedding similarity space. This will implicitly correspond to sampling low-RMSD values with high confidence in the predictions made, as also shown by the strong (anti-)correlation with the pLDDT.

### 2. A reliable and robust generative procedure can be achieved by an appropriate choice of effective temperature

When using MC sampling as a generative procedure, higher temperatures correspond to sampling regions of higher average energy. By contrast, lower temperatures constrain exploration to a narrow basin around the reference sequence.

The mathematical convergence properties of the Markov chain associated with the MC trajectory guarantee that the generated mutants will be distributed around a well-defined energy region, once equilibrium is reached. This can be readily observed in Fig. 6, where we show the energy as a function of the number of MC steps for different representative runs at different temperatures. As the number of MC steps (i.e., attempted mutations) increases, the energy plateaus around a value that increases as the sampling temperature increases. In all the cases analysed, if we choose a temperature for which the energy plateaus to a value below *E* ≈ 0.005, our algorithm reliably generates mutants with very low RMSD of the protected residues, below 1Å, as well as with high prediction confidence, with most mutants showing an average pLDDT above 0.7.

**Figure 6.**
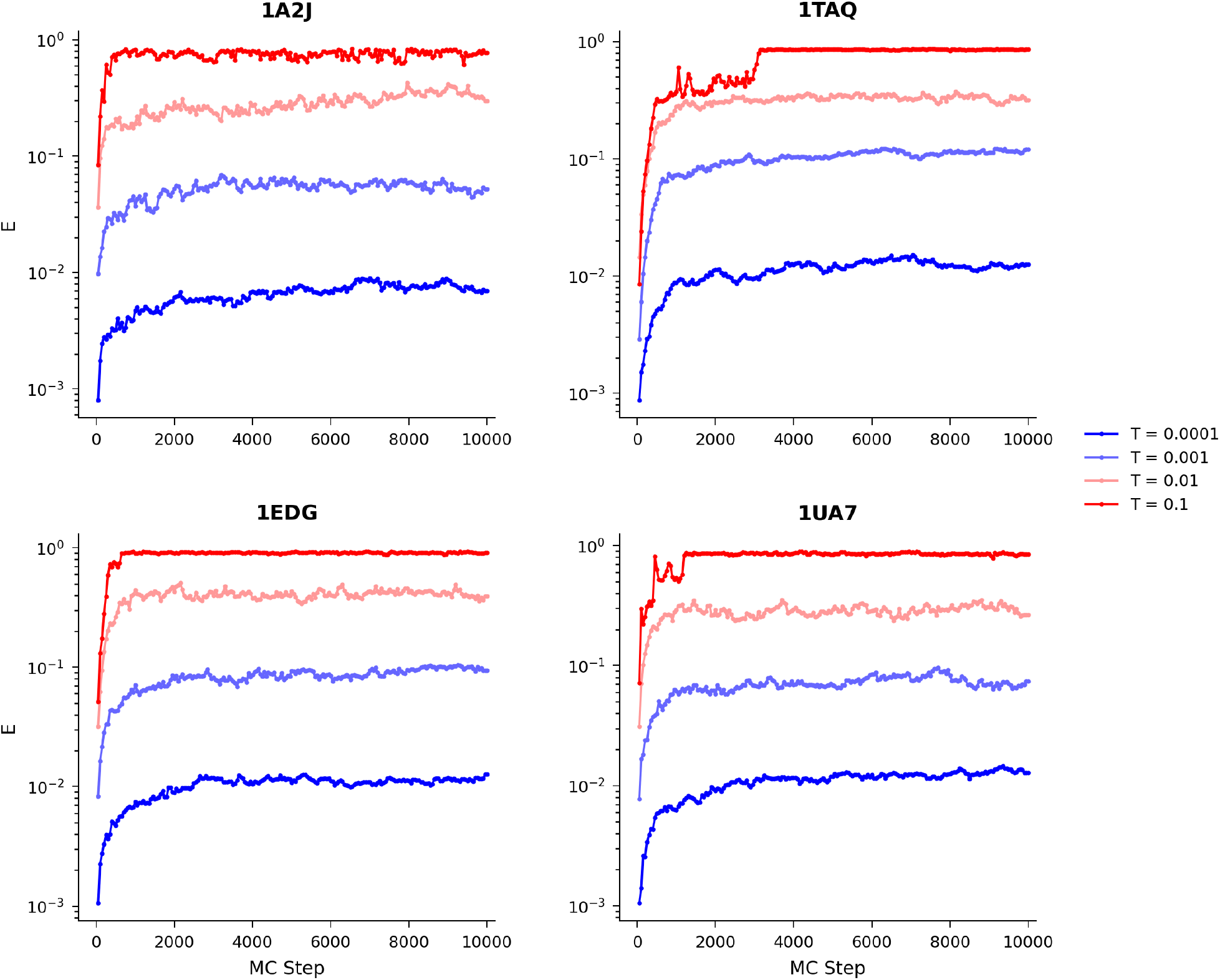
Trajectories of energy vs. number of MC steps during the generative procedure for the four representative enzymes from Fig. 3. Low to high sampling temperatures are presented as a colour gradient from blue to red, respectively. After some initial transient, it is clear that the trajectory plateaus around an equilibrium energy value that is a monotonic function of the temperature, as expected from the properties of MC sampling.

An alternative representation of the same data can be seen in Fig. 7, where we report the probability density function (PDF) of the mutant energies and protected residue RMSDs. Even in this case, it is clear that the distribution of mutants peaks both in energy and RMSD space, with the mode of the distribution moving to larger values for larger temperatures. Curiously, at high enough temperatures we also observe a phenomenon which we can relate to melting. For very low and very large temperatures, the algorithm produces distributions a single minimum; however, around a critical temperature the system shows a bimodal distribution with peaks both at high and low RMSD, corresponding to an equilibrium between what we can interpret as an ordered (low RMSD) and liquid (high RMSD) phase. Anecdotally, but somewhat interestingly, our simulations also demonstrated that starting from a sequence generated at very high temperature, the system is usually unable to find the minimum even after reverting the temperature below the critical value, at least not given sampling of a few thousands of MC steps. A similar situation occurs when one quenches a fluid via molecular simulations, where the system can remain trapped in a disordered, long-lived amorphous state that is not representative of the equilibrium configuration of that temperature. In practical terms, the presence of this hysteresis means that for the generative procedure to be successful, one should always start and remain at a temperature below the critical melting temperature. While we empirically observe that a temperature of below 0.001 always satisfies this condition for all enzymes we have simulated, this temperature should be expected to be system-dependent. Furthermore, although we do not study this effect here in detail, we generally expect this temperature to also depend on the specific pLM used. Determining this critical temperature is relatively easy, based on the fact that above this threshold, very unfavourable mutations are accepted, resulting in random structures being rapidly generated which completely lose any resemblance to the reference enzyme. Based on previous observations, a simple protocol to find the maximum sampling temperature for any system involves running a series of simulations at logarithmically spaced temperatures and monitoring the equilibrium energy in each case. For a small subset of the samples generated, the corresponding RMSD of the protected residues can be measured. One can then simply proceed with the generative procedure at or below the highest temperature at which the RMSD is below an acceptable level.

**Figure 7.**
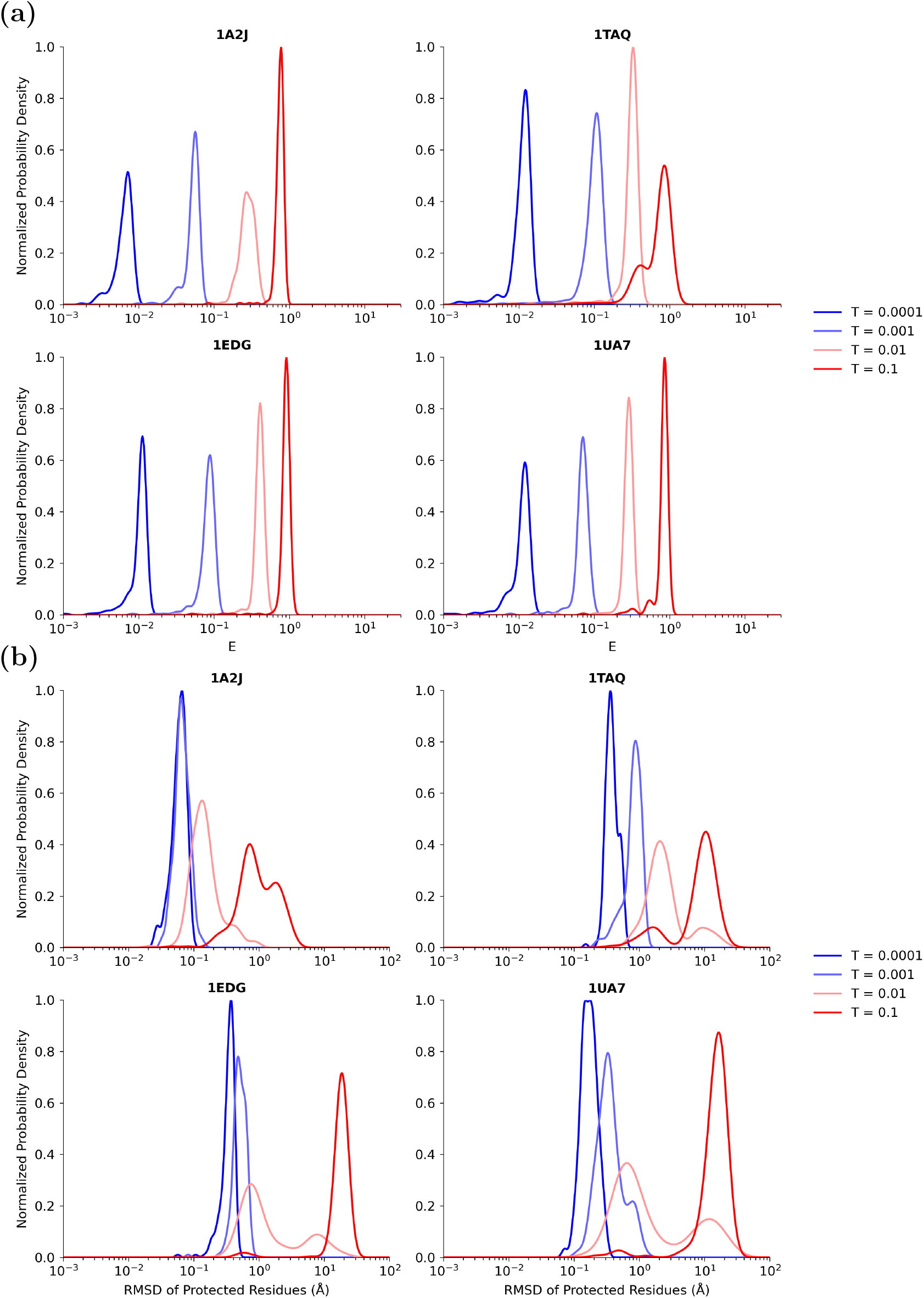
Temperature-dependent distributions of energy and RMSD across the four representative enzymes from Fig. 3. a) Probability density function (PDF) of the energy values observed (corresponding to the energy of different mutants) as a function of sampling temperature. b) PDF of the RMSD values as a function of sampling temperature. As a result of the generative procedure being based on a MC approach, sampling at increasing temperatures is equivalent to shifting the average value of the energy and RMSD sampled to higher values, providing an easy and physically justifiable way to control the generative procedure. The PDF is estimated using Kernel Density Estimation and Scott’s rule^26^ for the choice of the Gaussian kernel width.

### 3. The sampling temperature only weakly constrains sequence identity

While higher sampling temperatures lead to increased structural deviations from the reference enzyme, our energy function does not explicitly bias towards sequences close to the original one. Instead, the generative process samples sequences according to their embedding-defined energy, treating all sequences with equivalent energy equally, regard-less of how similar they are to the reference. As a result, even mutants with low sequence similarity may be sampled at low temperatures, provided they preserve the local embedding environment of protected residues.

This decoupling of structure and sequence is shown in Fig. 8, where for the previously reported enzymes, we present the distribution of sequence identity for the generated mutants; here, defined as the fraction of non-mutated residues compared to the reference sequence. As expected, higher temperatures broaden the distribution, enabling more divergent sequences. However, even at the lowest temperature tested (*T* = 10^*-*4^), where average protected residue RMSDs remain well below 1Å, sequence identity can drop to as low as 20%, corresponding to hundreds of residue differences.

**Figure 8.**
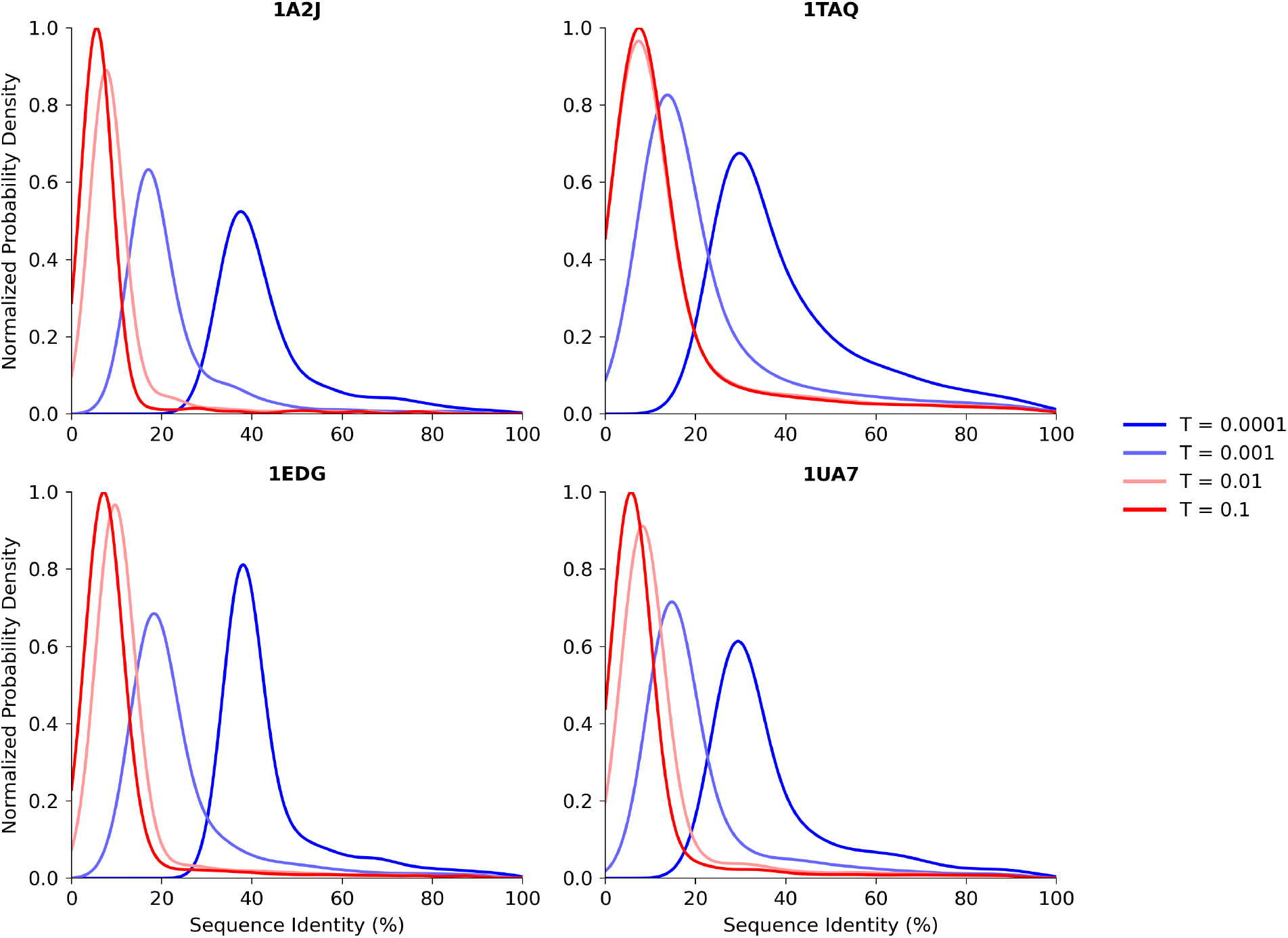
Probability Density Functions (PDF) of the sequence identities obtained for the four representative enzymes from Fig. 3 at different temperatures. Although in principle our algorithm does not explicitly bias sequence similarity, reducing the sampling temperature clearly results in structures of higher similarity. For the enzymes presented here, a sequence identity of 40% involves the mutations of approximately 110 amino acids for 1A2J and 500 amino acids for 1TAQ. Similarly to Fig. 7, the PDF is estimated using Kernel Density Estimation and Scott’s rule^26^ for the choice of the Gaussian kernel width.

These results highlight how our algorithm can simultaneously maintain local structural integrity and achieve substantial sequence diversity; a property that is essential for effective exploration of sequence space in enzyme engineering, as it enables the generation of highly novel variants while avoiding mutations that would compromise functionally critical regions. Importantly, the ability to generate diverse sequences without compromising structural constraints avoids a common failure mode of AI-driven generative models: mode collapse, where a model fails to produce diverse samples, and instead, yields a few high-probability outputs.

Taken together with the strong correlation between embedding energy and RMSD, these findings confirm that sampling low-energy regions is sufficient to ensure structural conservation at protected sites, while still supporting broad sequence exploration. This balance between constraint and diversity positions our method as a powerful tool for designing enzyme variants with high functional potential.

## III. DISCUSSION

The method presented here introduces a general and interpretable framework for enzyme sequence generation that directly addresses several limitations of existing approaches. Below, we discuss the key advantages of our method relative to prior generative models, its current limitations, and the broader implications for protein design.

Our approach distinguishes itself from existing models in two fundamental ways. First, it enables explicit preservation of user-defined structural motifs, such as catalytic sites. This ability is similar to that of complex multi-step pipelines, including those with backbone diffusion, sequence in-painting, and refolding;^27,28^ but unlike these end-to-end deep learning models, our method guarantees preservation of key residues by design. This means we can guarantee that any engineered mutant will retain the essential catalytic site, a feature typically absent in simpler, one-shot generative approaches.^11,12^ Second, a significant advantage of our model is that it doesn’t require any additional training beyond the initial pre-trained pLM — a stark contrast to all other current models. This latter aspect offers two additional advantages.

First, the computational cost of our generative procedure is minimal, determined solely by the pLM’s sequence-to-embedding transformation. This step is easily handled by low-end computational hardware, including desktop machines. Our current basic Metropolis MC implementation already enables the generation of hundreds of diverse, structurally constrained mutants within minutes on a single GPU. While equilibration and generation can be significantly sped up through the use of more complex algorithms (e.g., parallel tempering) or by simply parallelising the mutation process and leveraging batch predictions with a Heat Bath acceptance method instead of Metropolis MC,^22^ the baseline performance is already practical for many use cases.

The second advantage is our model’s generality. This stems from its pure transfer learning approach, which only requires the pLM to generate expressive, context-dependent embeddings that capture information about each residue’s environment. Because we solely extract information from the pLM (RMSD calculations are only for validation of our hypothesis, not generation), we expect our model to provide optimal mutants even for enzymes and proteins with unknown structures. This makes the approach particularly attractive for working with large databases of uncharacterised sequences lacking structural annotations. In principle, by not relying on any additional fine-tuning, our model is compatible with any modern pLM that provides context-aware embeddings. While we use ESM-2 due to its proven structural relevance (e.g., as used in ESMFold), alternative models such as aminoBERT^8^ or the more recent AMPLIFY^10^ could also be employed.

Beyond practical advantages, our method benefits from well-established theoretical guarantees. By employing a well-established, decades-old MC sampling method, we benefit from its mathematically provable convergence. This delivers robust sampling, ensuring that our algorithm generates low-energy mutants with exponentially higher probability as the process unfolds, regardless of their distance in sequence space from the original. This capability not only maximises diversity but also effectively avoids common pitfalls like mode collapse that can plague purely deep neural network-based generative approaches. Additionally, and perhaps most powerfully, our procedure frames the generative process as sampling of an energy landscape. This unique formulation allows for systematic and flexible biasing of the search towards sequences with user-defined properties. Any feature predictable from a sequence, for instance, using a deep neural network, can be incorporated into the energy function as an auxiliary term. This lets us penalise undesired deviations and optimise toward specific target values for features like an enzymes’ ease of expression, or its thermal stability. Crucially, the MC framework imposes no differentiability or continuity constraints on these predictive models, enabling seamless integration of both regression and classification outputs. Unlike one-shot generative approaches where tuning the relative influence of multiple objectives is often intractable, our method allows for explicit and interpretable weighting of each energy component.

This inherent modularity and tunability makes our approach exceptionally well-suited for multi-objective enzyme design, where diverse and often competing constraints must be balanced with precision.

## IV. CONCLUSIONS

In conclusion, we integrate a pLM with a statistical mechanics-based approach to introduce a generative algorithm that is able to create an optimal pool of mutants for experimental screening, starting from any initial enzyme. By construction, these mutants can be almost arbitrarily different in their sequence from the original one, yet preserve the catalytic site and its local structure, and are thereby depleted of deleterious mutations that would likely result in a loss of catalytic activity.

A core strength of the method lies in its theoretical foundation. By basing our generative procedure on solid statistical mechanics concepts and using a well-established algorithm such as MC sampling, we provide a clear and robust interpretation of our generative procedure, its convergence properties, and the role of the effective temperature as an intuitive control parameter. In addition to increasing our ability to understand and control the results, this hybrid method can be used to bias the generative procedure toward enzyme sequences that display any feature for which a sequence-feature function exists, simply by adding this feature as an additional term in the energy function. In the future, this aspect will allow us to go beyond the simple preservation of catalytic site shown here, and directly optimise for other application-relevant quantities, e.g., thermal stability, solubility or toxicity.

Finally, while our exposition focused on a limited set of four enzymes, we emphasise that the same qualitative trends and conclusions are held across a broader benchmark of 13 structurally and diverse enzymes, spanning lengths from approximately 100 to 800 residues with extremely low sequence similarity. For each case, our algorithm reliably generated over 12,500 mutants with catalytic site RMSD smaller than 2Å. This library is provided in the Supplementary Information.

Overall, our results represent a step forward in developing interpretable and controllable generative algorithms for enzyme engineering and, more broadly, protein design. By coupling the power of pLMs with the transparency of classical simulation techniques, our framework offers an alternative to current one-shot generative approaches. This hybrid strategy enables targeted exploration of sequence space with explicit control over structural constraints—crucial for optimising enzyme function through directed evolution. We suggest that the prevailing emphasis on end-to-end sequence generation may be misplaced for many practical applications, where interpretability, modularity, and precise constraint enforcement are more valuable than automated novelty. In this context, our approach offers a compelling path toward more rational and efficient protein engineering.

## V. METHODS

### 1. Folding via ESMFold and AlphaFold2

To validate our results and ensure robustness, we employed two state-of-the-art protein structure prediction tools: ESMFold and the ColabFold implementation of AlphaFold2,^29^ to model the structures of the generated sequences and compare them to their respective reference enzymes. This dual-model approach helps guard against adversarial artifacts specific to a given predictor and strengthens confidence in the structural plausibility of the designed variants.

For ESMFold, calculations were run using Nvidia T4 GPUs using AWS, and took about 30 seconds per structure. Regarding ColabFold, the latter was used with the following parameters: template usage was disabled (`use templates=false`) to prevent bias from known structures. Multiple sequence alignments (MSAs) were generated using MMseqs2^30^ against the UniRef sequence databases^31^ (`msa mode=mmseqs2 uniref env`). The model type utilised was `alphafold2 ptm`, which incorporates predicted TM-score and alignment error metrics. Five distinct models were generated per sequence (`num models=5`), each undergoing three recycling iterations (`num recycles=3`) to refine predictions. Amber relaxation was not applied (`num relax=0`) to expedite computation. All ColabFold predictions were executed on NVIDIA A100 GPUs via Google Colab, with an average runtime of approximately 1500 seconds per structure. For our calculations, we used the structure with the highest overall confidence, as measured by the pLDDT. The RMSD between the predicted structures of the original enzymes and their existing experimental structures (codes 1A2J, 1EDG, 1TAQ, 1UA7 in the PDB database) is reported in Table S-1, along confidence metrics of the predictions.

### 2. RMSD calculations

RMSD between protein structures is defined up to a global translation and rotation. In all cases, we apply the optimal superposition that minimizes the RMSD, computed using the Kabsch algorithm.^32^ For the analyses presented in the main text, the alignment is restricted to the protected region of the enzyme; that is, the subset of residues whose geometry we aim to conserve, rather than the entire structure. To ensure a rigorous comparison, RMSD for the protected (immutable) residues is calculated using all heavy atoms, including side chains. For the remaining residues, only backbone atoms (C, C_*α*_, O, and N) are considered, as these atoms are universally present across all amino acid types and thus allow for consistent structural comparison.

## Acknowledgments

We kindly acknowledge Dr Daniele Visco for general discussions about the methods used and a critical feedback on an initial version of the manuscript. Q.T and S.A-U acknowledge computational resources and support provided by the Imperial College Research Computing Service (http://doi.org/10.14469/hpc/2232). They are grateful to the UK Materials and Molecular Modelling Hub for computational resources, which is partially funded by EPSRC (EP/T022213/1, EP/W032260/1 and EP/P020194/1).

## Code and data availability

The raw data for the different enzymes generated is available at the following URL https://doi.org/10.5281/zenodo.15696797. The code used to implement the simulations to produce the data is fully open source and under an MIT licence, and will be released soon on GitHub.

## Authors’ contributions

S.A-U conceived the generative algorithm, analysed the data and wrote the manuscript. A.W, Q.T and A.R wrote the code used to generate the data, performed the numerical simulations and analysed the data. J.L analysed the data. S.A.-U, A.W., A.R and J.L. wrote and revised the manuscript.

## Notes

https://doi.org/10.5281/zenodo.15696797

### Competing Interest Statement

A.W., A.R., and S.A.-U. are affiliated with AminoAnalytica, a computational biology startup. The remaining authors declare no competing interests.

### Summary of Updates

This version of the manuscript has been revised to update the following: author affiliations updated

