## Supplementary Information for "Mind the Gap: An Embedding Guide to Safely Travel in Sequence Space"

#### 1. A comparison between ESMFold vs. AlphaFold2

Figure S-1 compares structural predictions from ESMFold and AlphaFold2 across 30 mutants for each of the four enzymes discussed in the main text. The right panel highlights the low-RMSD regime in the left panel. In most cases, low-energy structures that are predicted to have a low RMSD with ESMFold are also confirmed to have a low RMSD by AlphaFold2.

As shown in Fig. S-2, structures with low predicted energy also exhibit high confidence scores (pLDDT) in both models, with a strong correlation evident from the clustering of points along the diagonal. This consistency further supports the reliability of low-energy, low-RMSD predictions; precisely the class of structures most relevant for practical applications in enzyme design, where preserving the geometry of the catalytic site is critical.

It is worth noting, however, that the data is slightly biased in favor of ESMFold, as ESMFold-derived embeddings were used during mutant selection. This may account for the marginally lower RMSD and higher pLDDT values observed in ESMFold predictions relative to AlphaFold2.

**Table S-1**. **Structural prediction accuracy between ESMFold and AlphaFold2 for the reference sequence of the enzymes discussed in the main text.** RMSD values are reported in Å, and pLDDT scores are presented between the range of [0,100], with 100 representing the highest confidence. RMSD and pLDDT assess the fidelity of predicted structures relative to experimentally determined conformations and the model confidence in the prediction, respectively. In 1TAQ, the high RMSD stems from the wrong predicted alignment of two domains, each of which, however, is internally described with very high accuracy.

| PDB Code | ESMFold | | AlphaFold2 | |
|---|---|---|---|---|
| | RMSD (Å) | pLDDT | RMSD (Å) | pLDDT |
| 1A2J | 2.59 | 94.1 | 2.76 | 96.1 |
| 1EDG | 1.42 | 93.1 | 1.47 | 97.0 |
| 1UA7 | 1.22 | 93.0 | 1.36 | 97.8 |
| 1TAQ | 30.3 | 87.4 | 33.7 | 90.5 |

#### 2. RMSD and pLDDT outside the protected region

Fig. S-3 compares the RMSD of the protected residues with that of the rest of the enzyme, while Fig. S-4 reports the confidence metric for these predictions. We see for both the protected and unprotected regions, lower sampling temperatures lead to lower distortions, while the local structure of the protected region is always better maintained, with markedly lower RMSD. This result is a feature of our procedure, which only biases sampling to guarantee conservation of the environment for those residues specifically defined by the users, while allowing the rest of the enzyme to freely sample structural diversity. At the same time, it should also be noted that confidence metrics such as pLDDT also invariably deteriorates outside the protected region. However, this is a reflection of the fact that the relatively high randomness of the sequence outside the protected region leads more naturally to random coils rather

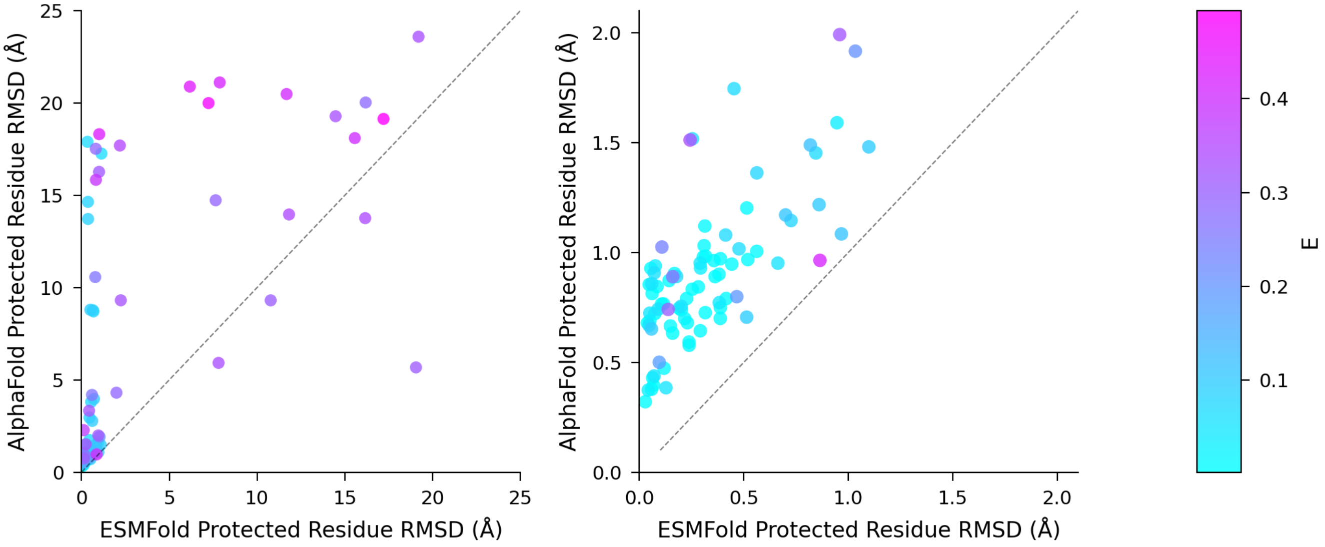


**Figure S-1**. **Correlation between ESMFold and AlphaFold2 RMSD prediction for a random subset of mutants generated during our simulations**. The subset was specifically chosen to sample a wide range of energies and RMSD of the active site. As somewhat expected, predictions are more highly correlated in the small RMSD region ($\leq 1.5$ Å), as shown in the inset on the right, which also generally corresponds to sequences of higher similarity with the original ones (as discussed in the main text). High-energy regions often correspond to almost random sequences, for which we do not expect a protein folding algorithm to work particularly well because of the lack of a reference system close enough for the inference process to provide a reliable result, as also shown by the drop in the confidence metric, see Fig. S-2. However, such high-RMSD sequences are also irrelevant from a practical point of view, considering keeping the active site intact is required to preserve catalytic activity.

than alpha helices or beta sheets, whose dynamical, unstructured nature is inherently associated to lower pLDDT values.

### 3. Additional library of variants generated in this study

To show the generality of our approach, we apply our generative procedure on a larger set of 13 different enzymes, covering different folds and families. This library of 12,500 sequences (all with RMSD smaller than 2Å) was curated and validated through structure prediction and is available at `https://doi.org/10.5281/zenodo.15696797` to support further experimental testing and integration into directed evolution pipelines. Selected results from this library are reported in Table S-2.

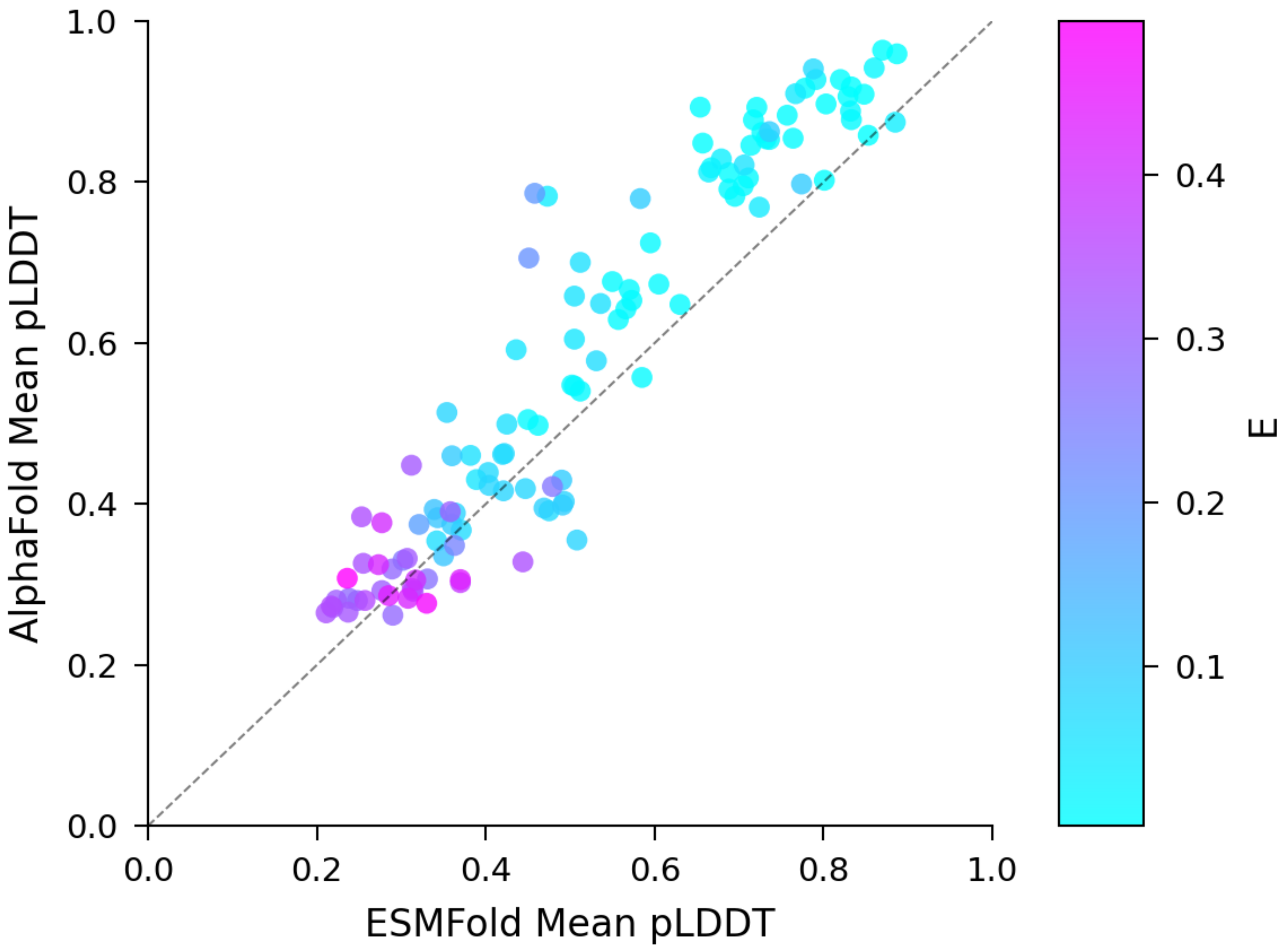


**Figure S-2**. **Correlation between ESMFold and AlphaFold2 pLDDT confidence scores for a random subset of mutants generated during our simulations**. There is a strong correlation between the pLDDT from both models, which is evident from the clustering of points along the diagonal. Predictions for lower energy mutants tend to have a higher confidence. The same subset from S-1 was used for this analysis.

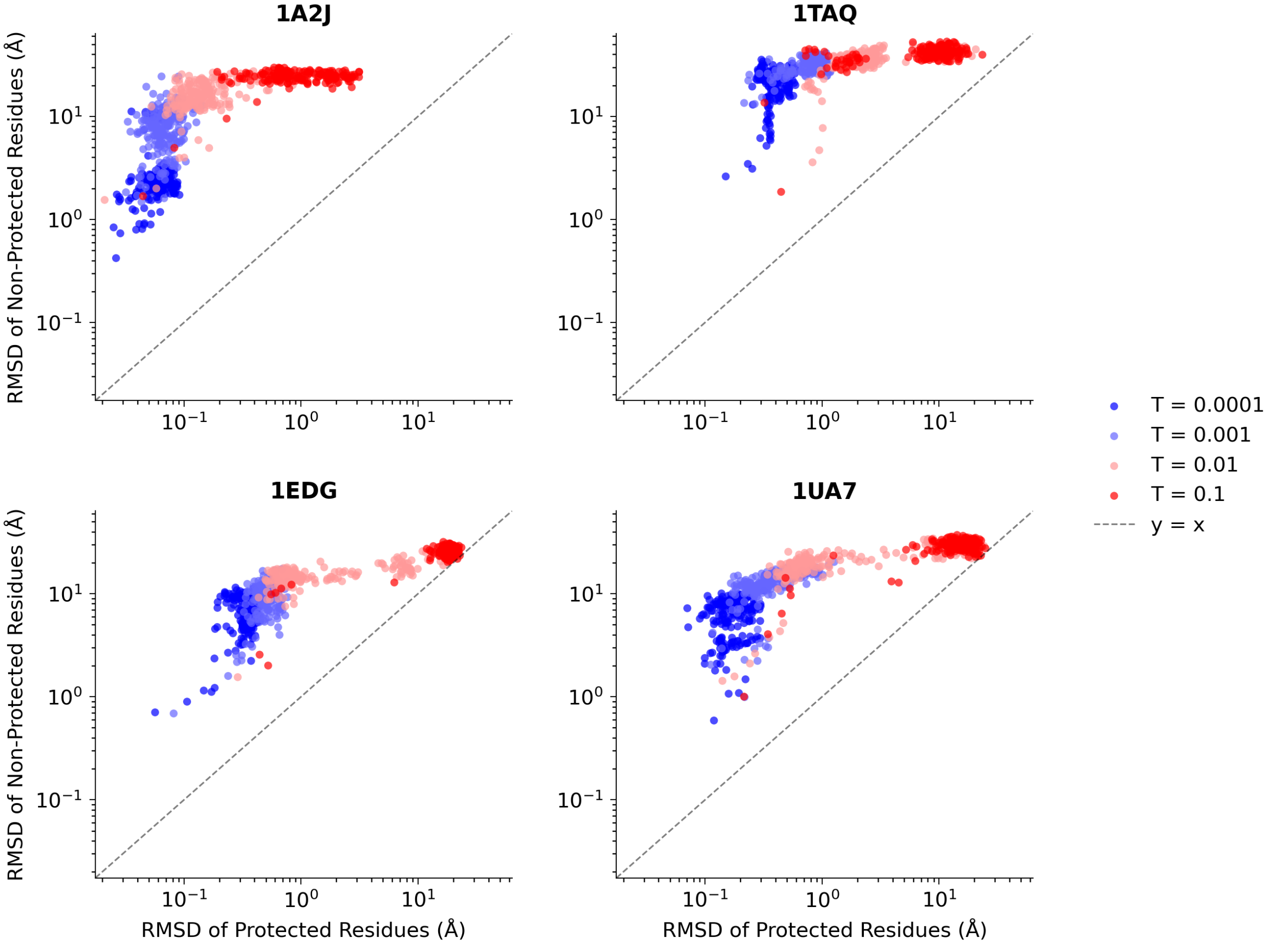


**Figure S-3**. **Comparison of RSMD for protected residues vs. non-protected residues for all four representative enzymes from Fig. 3, as a function of sampling temperature.** In general, lower temperatures correspond to lower RMSD values for the entire enzyme. However, the local structure of the protected region is always lower due to the residue preservation nature of our generative method.

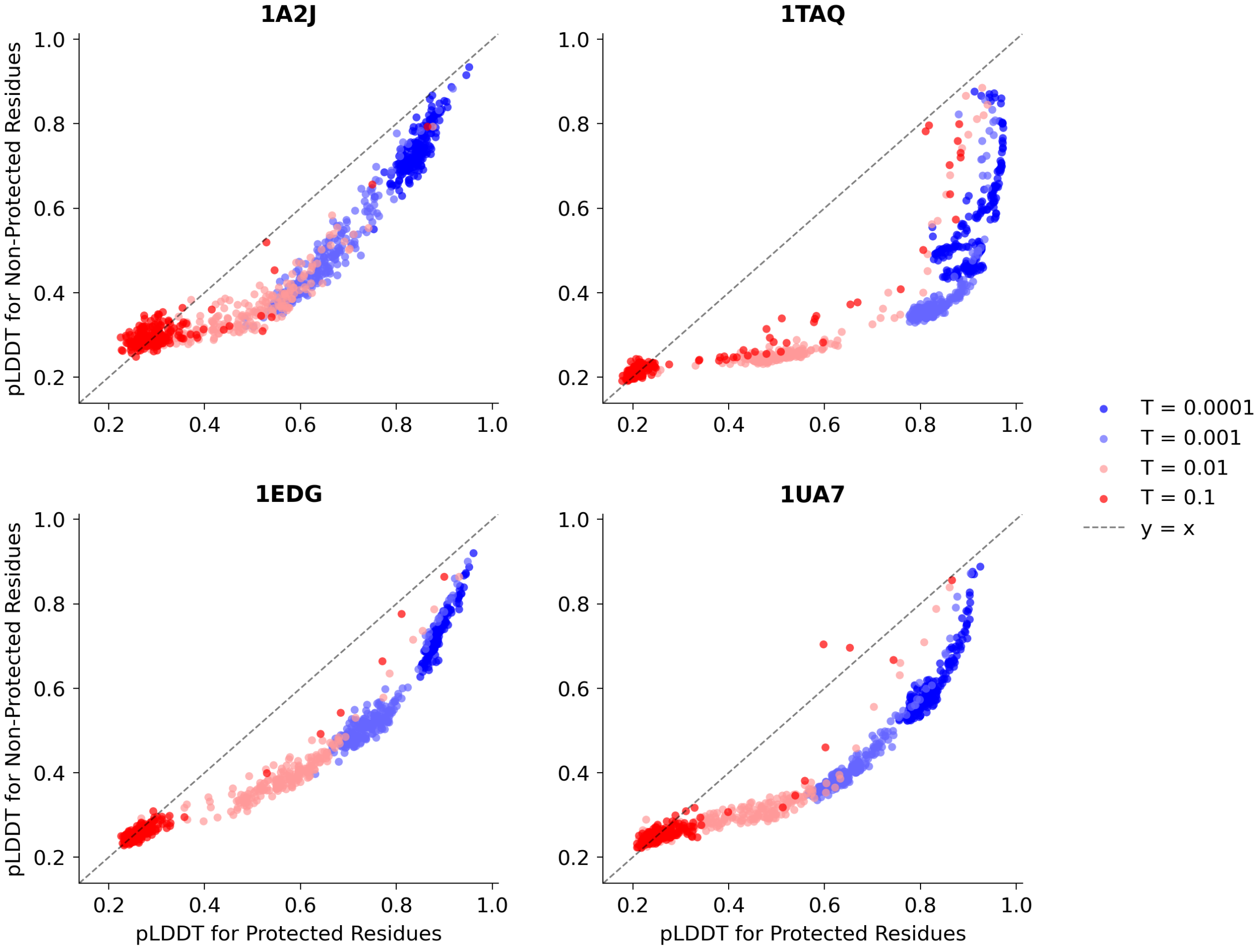


**Figure S-4**. **Comparison of pLDDT for protected residues vs. non-protected residues for all four representative enzymes from Fig. 3, as a function of sampling temperature.** As discussed in the main text, lower temperatures result in higher pLDDT values, with a slight deterioration for non-protected residues due to randomly occurring structural shifts from more confident alpha helices or beta sheets to less confident random coils.

**Table S-2**. **Structural conservation of the active site residues from a sample of mutants generated using our MC sampling approach.** For 13 representative enzymes (4 in the main text plus an additional 9), the table reports the protected site residues, the root-mean-square deviation (RMSD) of those residues between the parent and three mutants (**S1**–**S3**), and the corresponding sequence similarities. Protected residues have been chosen to represent those corresponding to the catalytic core of the enzyme. In all systems, the structural deviation in the catalytic core remains extremely small, with a mean RMSD of this sample of 39 enzymes of 0.355Å, despite having sequence identities as low as 2%. These data illustrate that our generative strategy can explore distant regions of sequence space while preserving the geometry of the active site, thereby enriching the pool of mutants that are immediately amenable to experimental screening.

| **PDB ID** | **Class** | **Protected Residues** | **Protected RMSD (Å)** | | | **Sequence Identity (%)** | | |
|---|---|---|---|---|---|---|---|---|
| | | | **S1** | **S2** | **S3** | **S1** | **S2** | **S3** |
| 1A2J | Oxidoreductase | C30, C33 | 0.021 | 0.270 | 0.137 | 79 | 2 | 5 |
| 1A58 | Isomerase | R66, F71, M72, Q74, G83, A112, N113, K114, Q122, F124, H132, L133, H137 | 0.119 | 0.434 | 0.446 | 95 | 19 | 20 |
| 1AEC | Hydrolase | C25, H162 | 0.012 | 0.184 | 0.217 | 92 | 6 | 6 |
| 1EDG | Cellulase | R79, H122, N169, E170, H254, Y256, E307 | 0.057 | 0.480 | 0.422 | 96 | 10 | 14 |
| 1TAQ | Taq polymerase | D610, I614, E615, F667, Y671, K663, R659 | 0.130 | 0.425 | 0.479 | 95 | 25 | 27 |
| 1TCA | Hydrolase | S105, D187, H224, T40, Q106 | 0.042 | 0.500 | 0.379 | 91 | 6 | 9 |
| 1THX | Oxidoreductase | W36, C37, G38, P39, C40 | 0.037 | 0.353 | 0.471 | 50 | 10 | 9 |
| 1UA7 | Hydrolase | D173, E205, D266 | 0.070 | 0.476 | 0.416 | 29 | 6 | 10 |
| 1ZG4 | Hydrolase | S45, K48, S105, E141, N145 | 0.020 | 0.483 | 0.476 | 98 | 17 | 19 |
| 2PPN | Isomerase | H25, Y26, F36, F46, W59, Y80, Y82, H87, F99 | 0.028 | 0.481 | 0.490 | 97 | 35 | 37 |
| 4M6K | Oxidoreductase | E31, R71, F35, N65 | 0.034 | 0.388 | 0.386 | 98 | 22 | 21 |
| 4RQR | Oxidoreductase | C26, C29 | 0.008 | 0.199 | 0.468 | 83 | 2 | 2 |
| 9PAP | Hydrolase | C25, H159, N175, Q19, W177 | 0.030 | 0.496 | 0.451 | 97 | 15 | 16 |
